# Perivascular spaces in Alzheimer’s disease are associated with inflammatory, stress-related, and hypertension biomarkers

**DOI:** 10.1101/2023.06.02.543504

**Authors:** Francesca Sibilia, Nasim Sheikh-Bahaei, Wendy J. Mack, Jeiran Choupan

## Abstract

Perivascular spaces (PVS) are fluid-filled spaces surrounding the brain vasculature. Literature suggests that PVS may play a significant role in aging and neurological disorders, including Alzheimer’s disease (AD).

Cortisol, a stress hormone, has been implicated in the development and progression of AD. Hypertension, a common condition in older adults, has been found to be a risk factor for AD. Hypertension may contribute to PVS enlargement, impairing the clearance of waste products from the brain and promoting neuroinflammation. This study aims to understand the potential interactions between PVS, cortisol, hypertension, and inflammation in the context of cognitive impairment.

Using MRI scans acquired at 1.5T, PVS were quantified in a cohort of 465 individuals with cognitive impairment. PVS was calculated in the basal ganglia and centrum semiovale using an automated segmentation approach. Levels of cortisol and angiotensin-converting enzyme (ACE) (an indicator of hypertension) were measured from plasma. Inflammatory biomarkers, such as cytokines and matrix metalloproteinases, were analyzed using advanced laboratory techniques.

Main effect and interaction analyses were performed to examine the associations between PVS severity, cortisol levels, hypertension, and inflammatory biomarkers.

In the centrum semiovale, higher levels of inflammation reduced cortisol associations with PVS volume fraction.

For ACE, an inverse association with PVS was seen only when interacting with TNFr2 (a transmembrane receptor of TNF). There was also a significant inverse main effect of TNFr2.

In the PVS basal ganglia, a significant positive association was found with TRAIL (a TNF receptor inducing apoptosis).

These findings show for the first time the intricate relationships between PVS structure and the levels of stress-related, hypertension, and inflammatory biomarkers. This research could potentially guide future studies regarding the underlying mechanisms of AD pathogenesis and the potential development of novel therapeutic strategies targeting these inflammation factors.

## 1. Introduction

Perivascular spaces (PVSs) are fluid-filled spaces in the brain surrounding blood vasculature (Doubal et al., 2010). Their function is to facilitate the drainage of cerebro-spinal fluid (CSF) between the interstitial space and the blood vessel space, allowing the outflow of waste-soluble proteins from the brain (Zong et al., 2020). PVSs are a key part of the glymphatic system, which are responsible for cleansing the brain of neurotoxins. In healthy populations, an increase in PVS number and volume is seen during normal aging (Lynch et al., 2022), especially in basal ganglia, centrum semiovale and hippocampus (Wardlaw et al., 2020). In clinical populations, studies have shown an association between dilated PVSs and several neurological diseases, including Huntington’s disease (Chan et al., 2021), cerebral small vessel disease (Brown et al., 2018), and Alzheimer’s disease (AD) (Boespflug et al., 2018).

Increased PVS volume and number can be caused by the alteration of CSF flow in the interstitial space (Barisano et al., 2022). Among environmental and external factors disrupting this balance, stress has been found to delay the hemodynamic response and neurovascular coupling that regulate cerebral flow of metabolites needed for cerebral function (Dunlop & Liston, 2018). Higher levels of stress were linked to hyperproduction of tau and amyloid, neuropathological changes typical of dementia and cognitive impairment (Ennis et al., 2017). The activation of the Hypothalamus-Pituitary-Adrenal (HPA) axis is one of the physiological mechanisms to stress response that leads to cortisol production and regulation in the body (Justice, 2018).

High cortisol levels can affect blood pressure (BP) and cause hypertension (Ouanes & Popp, 2019). High BP induces a change in vessel dynamics that reduces perivascular pumping, and decreases the net flow of CSF in PVSs, with a consequent reduction of parenchymal waste transport (Ennis et al., 2017). Blood pressure is regulated by the renin-angiotensin-adrenal system (RAS). Angiotensin-converting enzyme (ACE) is an enzyme that is part of RAS and is involved in the production of Angiotensin II; high levels of Angiotensin II cause arterial stiffness and structural remodeling, which lead to hypertension (Popp et al., 2015) and enlarged PVSs (Mestre et al., 2017). ACE was found to accumulate in PVS of patients with cerebral amyloid angiopathy (CAA) and correlated with parenchymal Aβ load in AD (Ouanes & Popp, 2019).

Together with the release of cortisol, the interaction of pro-inflammatory factors with the RAS is also crucial for the maintenance of brain vasculature (Xue et al., 2020). In previous studies, ACE has been shown to exert a proinflammatory action on the endothelial and vascular smooth muscle cells (Dandona et al., 2007) by releasing tumor necrosis factor (TNF), interleukins, and matrix metalloproteinases (MMPs), that are involved in cell apoptosis and neurodegeneration (Benigni et al., 2010) (Sproston & Ashworth, 2018). This contributes to the onset of cerebral small vessel diseases by damaging endothelial cells and disrupting blood brain barrier (BBB) (**Figure 1**). MMP-2 and MMP-9 were found to be expressed more around astrocytes in Alzheimer’s disease, promoting accumulation of amyloid beta (Aβ) plaques. MMP-2 and MMP-9 expression is regulated by TNF-*a* and were found to be related to hypertension (Cancemi et al., 2020). Therefore, the action of tumor necrosis factor alpha (TNF-*a*), TNF-receptor-2 (TNFr2) and TNF-receptor-apoptosis-inducing-ligand (TRAIL), interleukin-6 receptor (IL-6r) (Blecharz-Lang et al., 2018) and C-reactive protein (CRP) (Hilal et al., 2018) play a role in the correct response of the immune system (Burgaletto et al., 2020).

**Figure 1:**
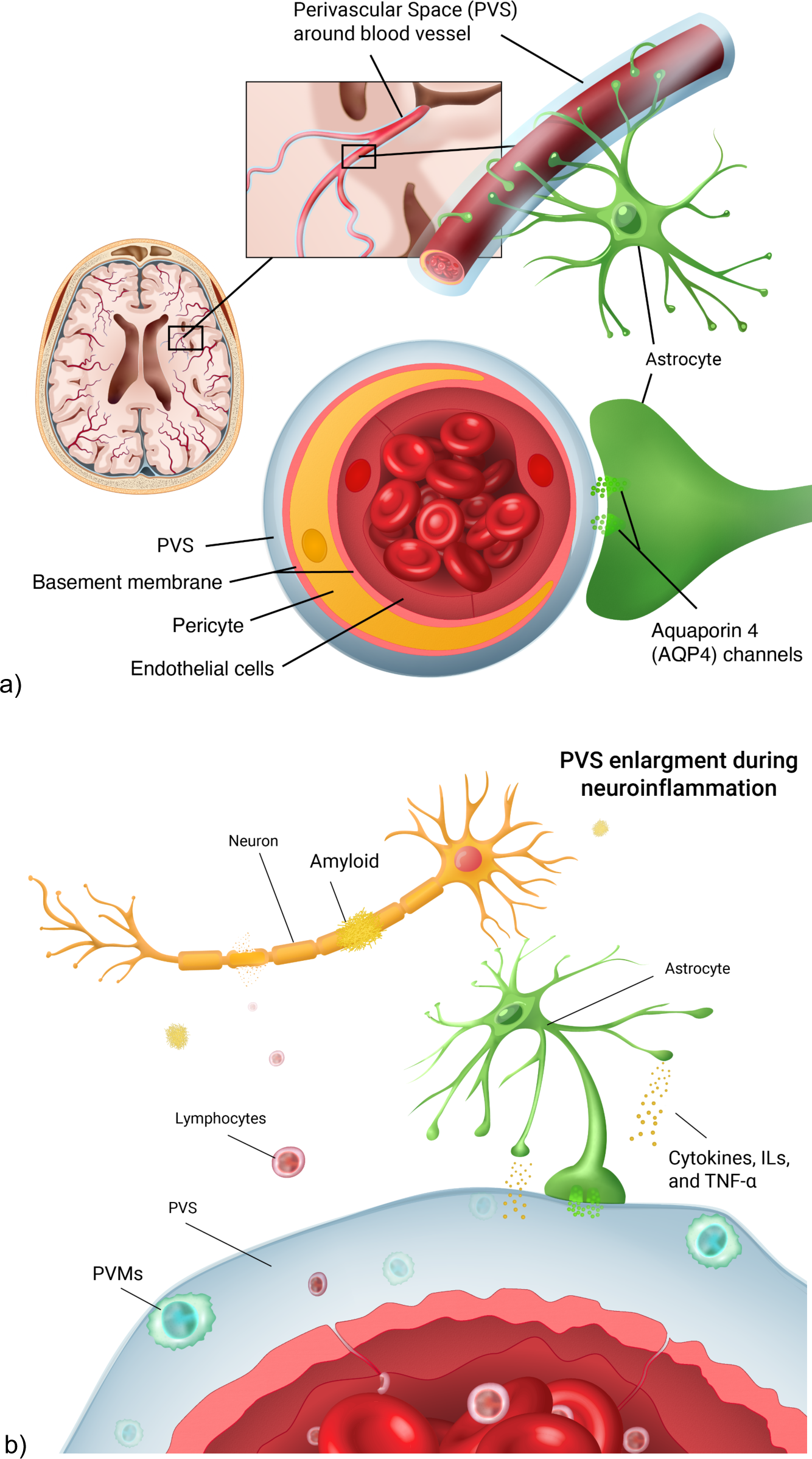
a) anatomical representation of the perivascular space (PVS) in the brain vasculature. PVS surrounds the blood vessel lumen. It is involved in the fluid exchange between blood vessels and brain parenchyma through the aquaporin 4 channels, located on the astrocyte endfeet. b) during neuroinflammation, lymphocytes cross the endothelial cell layer as a result of blood brain barrier leakage Perivascular macrophages (PVMs) then form, leading to PVS enlargements. As a result, there is a higher accumulation of neurotoxins such as amyloid beta, leading to neural death.

A perturbation in the physiological level of biomarkers in the brain can lead to alterations of the clearance system, where PVS is a crucial component. Therefore, exploring the relationship between alterations in PVS volume and plasma biomarkers of inflammation may help understand the mechanisms of neuroinflammation that contribute to the onset of cognitive impairment and neurodegenerative diseases which are understudied so far.

The objective of this study is to investigate the associations of plasma biomarkers of stress, hypertension and inflammation with perivascular space (PVS) volume fraction in older participants of the Alzheimer’s Disease Neuroimaging Initiative 1 (ADNI-1) cohort. We tested the main effect of each biomarker with PVS. We also tested whether PVS associations with cortisol and ACE are modified by levels of inflammatory biomarkers (TNF-*a*, TNFr2, TRAIL, CRP, IL-6, MMP-2 and MMP-9).

To our knowledge, this is the first study evaluating associations of stress-related, hypertension and inflammatory plasma biomarkers with PVS volume fraction in an elderly population, using a novel automated segmentation technique to measure PVS.

## 2. Methods

### 2.1. Participants

Participants were older adults from the ADNI-1 population (n=465, age range=55-90) using data from the 12-month follow-up after recruitment. Inclusion and exclusion criteria can be found on the ADNI dataset manual (https://adni.loni.usc.edu/wp-content/uploads/2010/09/ADNI_GeneralProceduresManual.pdf). In brief, ADNI inclusion criteria included age between 55-90, not enrolled in other studies, generally healthy, fluency in English/Spanish and Geriatric Depression Scale less than 6. Exclusion criteria included the use of specific medications within 4 weeks of screening, such as antidepressants, narcotic analgesics, and anti-Parkinsonian medications with anti-cholinergic activity.

In the current study, participants were excluded if they were missing data on demographics, medication history, physiological biomarkers or missing T1w-MPRAGE scans at month-12. From an initial number of 697 that had MRI scans acquired at 1.5T, there were 232 cases with missing biomarker information; therefore, our final sample size was n=465.

The selection of participants for this analysis included those who had complete demographic and plasma biomarker information at the 12-month assessment. Demographic, biomarker and imaging data were downloaded from the ADNI website (http://adni.loni.usc.edu) and are detailed in **Table 1**. Participants were categorized based on their diagnosis at ADNI enrollment into cognitively intact healthy controls (CN), persons with mild cognitive impairment (MCI) and persons with more advanced Alzheimer’s diseases (AD). Body mass index (BMI) was considered a hypertension-related factor, expressed as weight (kg) divided by height (m^2^) (for those participants expressing weight/height in lb/inch, we converted them in kg/m). Education was indicated in years; the number of APOE-4 allele copies was included as a marker of genetic predisposition for AD. Medications were dummy coded based on the type of medication participants reported using (ACE-inhibitors, steroids, other hypertension-related medication).

**Table 1:** Demographic information about ADNI-1 participants. Six people took more than one type of medication

| **ADNI-1 (m12), N = 465** |  |
| --- | --- |
| Sex |  |
| Male | 287 (61.7%) |
| Female | 178 (38.3%) |
| Age (mean ± SD) | 74.85 ± 7.09 |
| Body Mass Index (BMI, kg/m<sup>2</sup>) |  |
| <24.9 | 194 |
| 25<29.9 | 199 |
| >30 | 72 |
| Diagnosis |  |
| AD | 92 |
| MCI | 325 |
| CN | 48 |
| Education (mean ± SD) | 15.63 ± 2.97 |
| Medication history (ever used) |  |
| 0 (no medication) | 408 |
| 1 (Ace-inhibitors) | 33 |
| 2 (steroids) | 21 |
| 3 (other hypertension meds) | 15 |
| APOE4 (copies of alleles) |  |
| 0 copy | 222 |
| 1 copy | 181 |
| 2 copies | 62 |

### 2.2. Plasma inflammatory and physiological biomarkers

The plasma biomarkers used in this study were obtained from the ADNI website, part of the Biomarkers Consortium Plasma Proteomics Project RBM multiplex data (https://adni.loni.usc.edu/wp-content/uploads/2010/11/BC_Plasma_Proteomics_Data_Primer.pdf). A 190-analyte multiplex immunoassay panel was developed on the Luminex xMAP platform by Rules-Based Medicine (RBM, Austin, TX). The panel, referred to as the human discovery map, contains plasma proteins previously reported in the literature to be altered as a result of cancer, cardiovascular disease, metabolic disorders or inflammation. The Luminex xMAP technology uses a flow-based laser apparatus to detect fluorescent polystyrene microspheres which are loaded with different ratios of two spectrally distinct fluorochromes. Using a ratio of the fluorochromes, up to 100 different beads can be generated such that each of them contains a unique color-coded signature. The beads are bonded with either ligand or antibodies and then standard sandwich assay formats are used to detect the analytes. The beads are read one at a time as they pass through a flow cell on the Luminex laser instrument using a dual laser system. One laser detects the color code and the other reports biomarker concentration. Each biomarker considered in this study was collected at the 12-month visit (expressed in ng/mL), to correlate with imaging data collected at the same visit.

### 2.3. Medication Use

Since both antihypertensive medications and steroids can influence the blood pressure and cortisol levels, the history of hypertension-related medication use was evaluated for each participant. In particular ACE-inhibitors were considered as hypertension-related drugs, as their mechanism of action can influence physiological ACE levels. In ADNI-1, each participant provided information on the type of medications ever taken, the dose, the morbidity condition for which it was prescribed, and if they were currently taking the medication at the time of the study visit. The medication was assigned to a specific category (ACE-inhibitors, steroid or other hypertension-related medications). A medication variable defined whether each participant ever used medication related to hypertension or cortisol levels, in particular ACE-inhibitors, steroids (that control cortisol levels), or other hypertension-related medication (0= ‘no medication’; 1= ‘ACE-inhibitors’, 2= ‘steroids’, 3= ‘other hypertension medications’). Secondly, a 3-level variable was applied to indicate whether participants were currently using these medications (0= ‘no medication history’, 1= ‘past use of medications; 2= ‘current use of medications’).

### 2.4. Imaging

#### 2.4.1. Data acquisition

T1w MPRAGE images (n=465) were acquired using scanners from GE Healthcare, Philips Medical Systems, or Siemens Medical Solutions at 1.5T (voxel resolution was 1.25×1.25×1.2 mm^3^), TR=2400 ms, TE= 3.6 msl, flip angle=8.0 degree, FOV=24 cm, slice thickness=1.2mm).

We further replicated the same analyses using a subsample from ADNI-1 with MRI scans acquired at 3T (8-channel coil, TR = 650 ms, TE = min full echo, flip-angle = 8°, slice thickness = 1.2 mm, resolution = 256 × 256 mm and FOV = 26 cm, voxel size=1×1×1.2 mm^3^), as well as neuroinflammation, stress-related and hypertension biomarkers (n=108). This was done to replicate the same results when using data acquired at higher magnetic field strength, although the sample size for the 3T data is much smaller.

#### 2.4.2. Data processing

PVS segmentation follows a previously published technique from our lab (Sepehrband et al., 2021, Sepehrband et al., 2019). This is the first time that this method was used to segment PVS from T1w scans at 1.5T.In brief, after data wrangling and preprocessing, PVSs were mapped from T1w images: A non-local mean filtering method was applied to denoise the MRI T1w images (Manjón et al., 2010) which is based on how a voxel of interest is similar to neighboring voxels based on their intensity values. A filtering patch with a radius of 1 voxel was applied to retain PVS voxels while removing the image noise at a single-voxel level (Manjón et al., 2010). The Rician noise of MRI scans was considered as the noise reference level for the filtering algorithm. After this, a Frangi filter was applied (Frangi et al., 1998) to detect tubular structures (in this case PVS) on the T1w at a voxel level using the Quantitative Imaging Toolkit (QIT) (Cabeen et al., 2018). A range of 0.1–5 voxels was chosen for this step, as it enhances the detection of vessel-like structures. The output of this step is a probabilistic map of vesselness (Frangi et al., 1998), which was thresholded to obtain a binary PVS mask. The threshold of 0.00001 was used based on expert opinion to capture true positives and true negatives.

Freesurfer (v.7.1.1) was run to perform image pre-processing (motion correction and image normalization and skull stripping) and to obtain brain volume and parcellation, by running the *recon-all* module on the Laboratory of Neuro Imaging (LONI) pipeline system (https://pipeline.loni.usc.edu). PVS volumes were extracted from both total white matter and basal ganglia (BG), as well as for brain regions based on the Freesurfer’s Desikan-Killiany-Tourville adult cortical parcellation atlas (Klein and Tourville, 2012). The centrum semiovale (CSO) was selected because this region has been historically used as the clinical ROI to assess PVS, and it was obtained by adding regions parcellated by Desikan-Killiany atlas. A list of regions forming the CSO is found in Table 1. Appendix A shows an overlay presentation of the CSO.

The PVS volume fraction was calculated by dividing the PVS volume by the total volume for both CSO and basal ganglia in each participant. The regions forming the CSO are listed in **Table A1,** and **Figure A1** (the RGB values indicated in the table are from the Freesurfer atlas). Total hippocampus volume was obtained by Freesurfer volumetric analysis and was used as an indicator of AD-related brain atrophy. A visual representation of the PVS segmentation is shown in **Figure A3**.

### 2.5. Statistical analysis

PVS volume fractions for both basal ganglia and CSO were used as a dependent variable (DV). The different classes of biomarkers were considered as independent variables (IV) in separate models. To properly model the positive, continuous skewed PVS data, we used generalized linear regression specifying a Gamma distribution and log link function. We first analyzed the main effect relationships between PVS volume fraction and stress levels (Cortisol), hypertension risk (ACE) and inflammation (CRP, IL-6, TNF-*a*, TNFr2, TRAIL, MMP-2 and MMP-9) biomarkers separately. Secondly, we investigated the interactions between physiological (cortisol, ACE) and inflammatory biomarkers. Distributions of the inflammatory biomarkers are reported in **Figure A2**. Continuous independent variables used in interaction analyses were centered around their median values. Sex, age, BMI, and hippocampal volume were included as covariates in all analyses. BMI was scaled using z-transformation to reduce distribution skewness. In sensitivity analyses, we also included an indicator variable for APOE4 positivity as a model covariate. Although we used hippocampal volume as a risk indicator for cognitive impairment, we also included cognitive diagnoses (AD, MCI, CN) as model covariates (reported in **Section 4** of the Results in Appendix). Medications were included as model covariate, by coding them as factor variable as described in 2.3. The main effect and interaction models with Gamma distribution are described by the mathematical formulas reported below - a) main effect, b) interaction with Cortisol and c) interaction with ACE. The statistical models were run considering separately the (*pvs_vf_*) of the centrum semiovale and basal ganglia as dependent variables. – Equation 1:

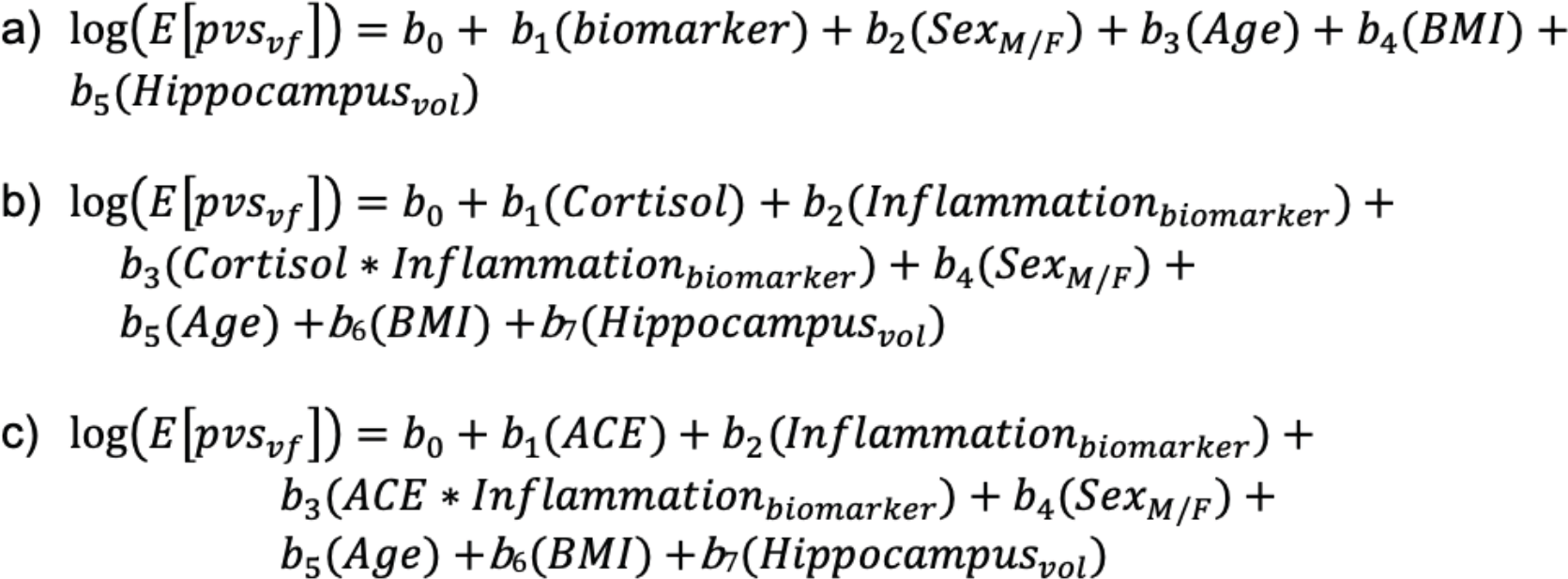

**Equation 1**: Regression models describing the main effect for PVS volume fraction. a) main effect of each biomarker; b) interaction between Cortisol and inflammatory markers; c) interaction between ACE and inflammatory markers. Regressions were run both for PVS centrum semiovale and basal ganglia

After removing the outliers for the PVS volume fraction (9 in total, calculated by multiplying 1.5 times the interquartile range) the final sample for the analysis is n = 456 participants. A two-sided p-value of 0.05 was used to determine statistical significance; regression residuals were investigated for normality. Statistical analyses were used RStudio (v.2022.07.2). We showed statistically significant fitted associations of either cortisol or ACE at two levels (minimum and maximum) of inflammatory markers.

## 3. Results

We report here from the 1.5T data; findings from 3T are included in the **Section 2** of the Appendix.

### 3.1. PVS in centrum semiovale (CSO-PVS)

#### 3.1.1. Main effects of Cortisol, ACE, MMPs

There was no significant association between PVS volume fraction in the CSO and cortisol, ACE or MMP levels. Beta coefficients, standard errors and p-values are shown in **Section 3** in the Appendix (**Figure A4**).

#### 3.1.2. Main effect of inflammatory biomarkers

Among the inflammatory biomarkers, there was no significant relationship with PVS volume fraction. CRP showed a tendency of significance (beta (SE) = 0.95 (0.03); p= 0.051), together with a significant effect of age (p<0.001) and BMI (p=0.003) on PVS volume fraction. When the medications were included in the model, the CRP association with PVS did not change. Other inflammatory biomarkers were not significantly associated with PVS in this region.

#### 3.1.3. Cortisol interaction analysis

In CSO-PVS, interactions of cortisol with inflammatory biomarkers showed inverse associations with PVS, such that higher levels of inflammation reduced cortisol associations with PVS; this was observed for TNF-α (interaction p=0.008), TRAIL (p= 0.013, t= -2.498), TNFr2 (p= 0.008), CRP (p=0.028) and MMP9 (p= 0.015) (**Figure 2**). Interactions of cortisol with MMP-2 and IL-6r were not statistically significant. Regression model results are shown in **Table 2**. Results were confirmed when information on whether participants ever used medications was added to the model.

**Figure 2:**
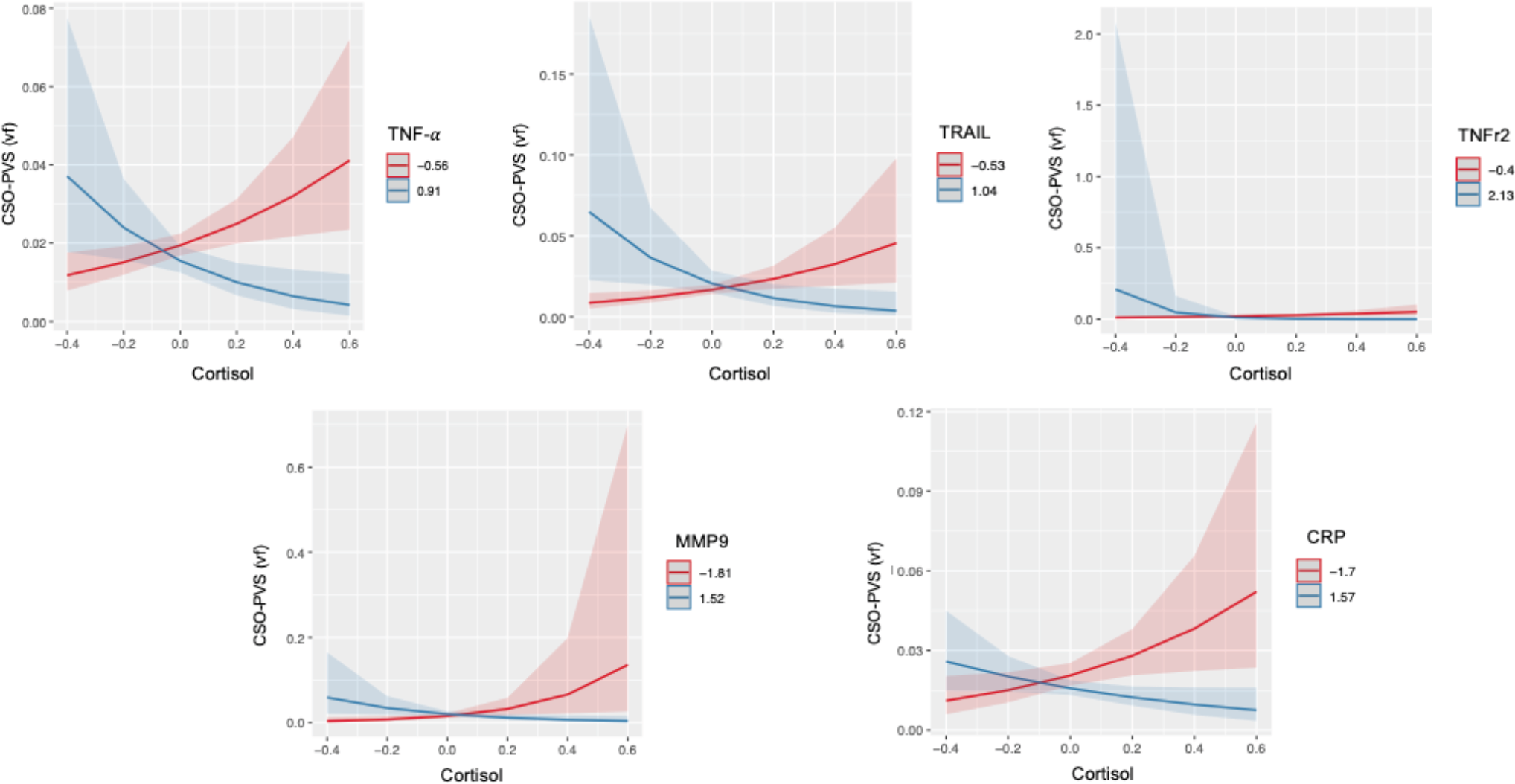
Statistically significant interactions between Cortisol and TNF-α, TRAIL, TNFr2, MMP-b9 and CRP in centrum semiovale (CSO) PVS volume fraction (vf). Plots indicate model-fitted cortisol-PVS curves at two different levels of each inflammatory biomarker. The two levels of inflammatory markers indicate the minimum and the maximum values. Shaded areas represent 95% confident intervals.

**Table 2:**
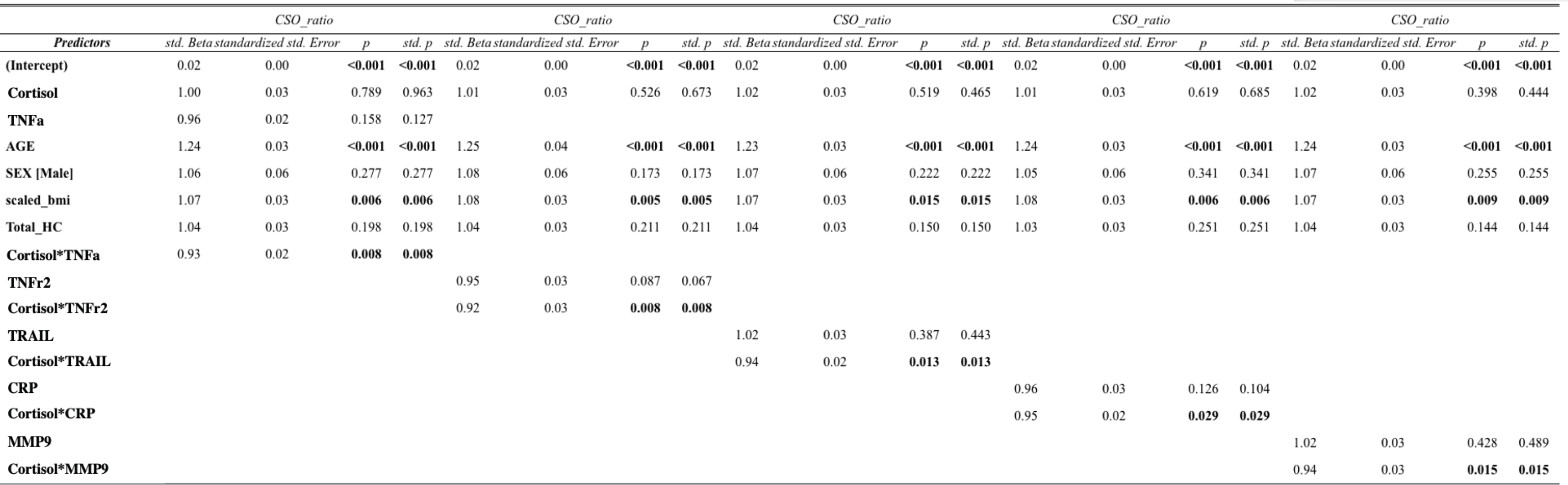
Beta coefficients, standard error and p-values for significant interactions between Cortisol and inflammatory biomarkers (TNF-α, TNFr2, TRAIL, CRP and MMP-9) in the centrum semiovale PVS volume fraction.

#### 3.1.4. Interactions with ACE

Interaction of ACE with TNFr2 showed inverse associations with PVS (**Table 3**), such that higher levels of inflammation reduced ACE associations with PVS volume fraction (p=0.006). There was also a significant inverse main effect of TNFr2 (p=0.014), indicating a positive association of ACE with PVS at the median values of TNFr2, and positive main effects of age (p<0.00001) and BMI (p= 0.003). The results were confirmed when data on whether participants ever used medications (interaction p= 0.006) and whether they were currently using medications (interaction p=0.006) were included in the model.

**Table 3:**
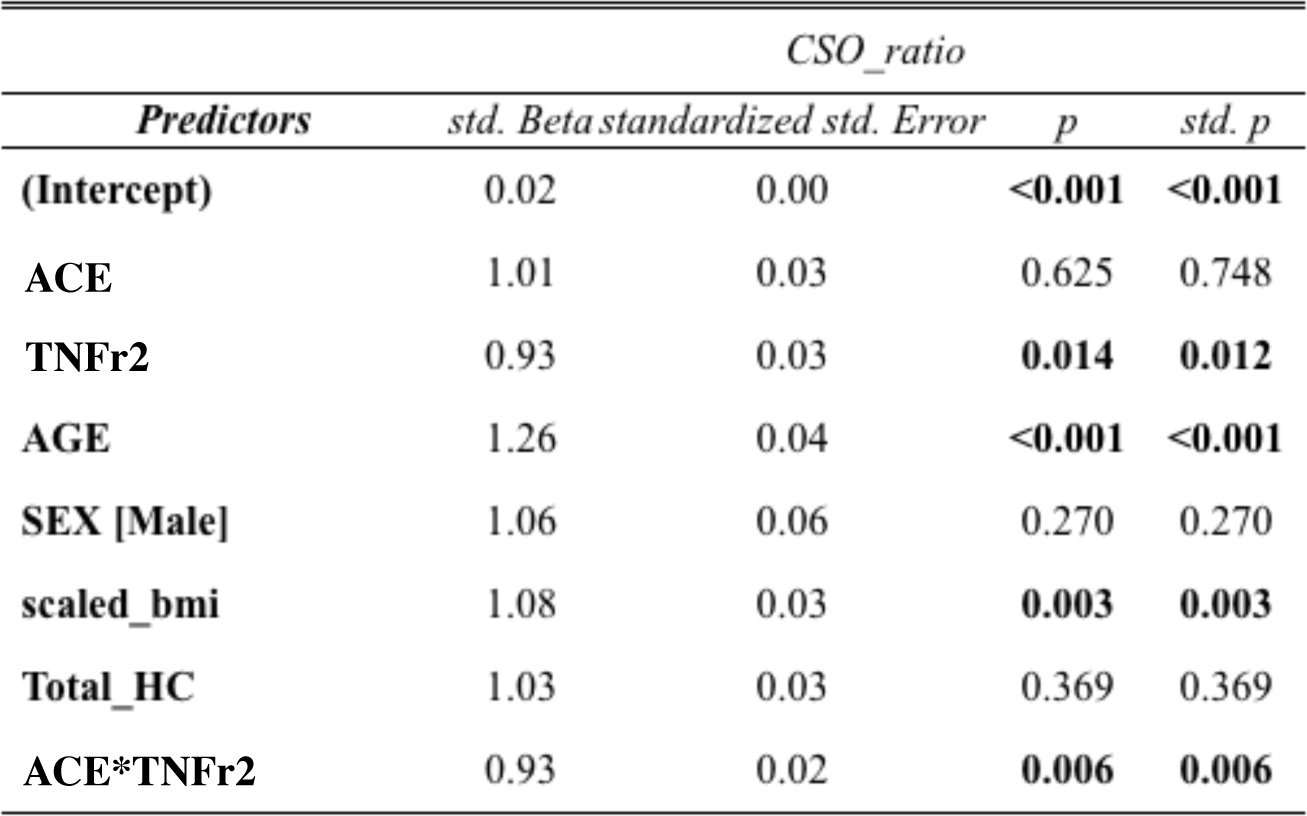
Beta coefficients, standard error and p-values for the significant interaction between ACE and TNFr2 in the centrum semiovale PVS volume fraction.

### 3.2. PVS in basal ganglia

#### 3.2.1. Main effects of Cortisol, ACE, MMPs

There were no significant associations between any of the physiological biomarkers and PVS volume fraction of basal ganglia. Regression model results are summarized in Table A4b in Appendix.

#### 3.2.2. Main effects of inflammatory biomarkers

A significant positive association between PVS in basal ganglia and TRAIL was seen (p= 0.049, **Table 4**). This result was confirmed when information on whether participants ever used medications (p= 0.049) and whether they were currently using them (p=0.035) were considered.

**Table 4:**
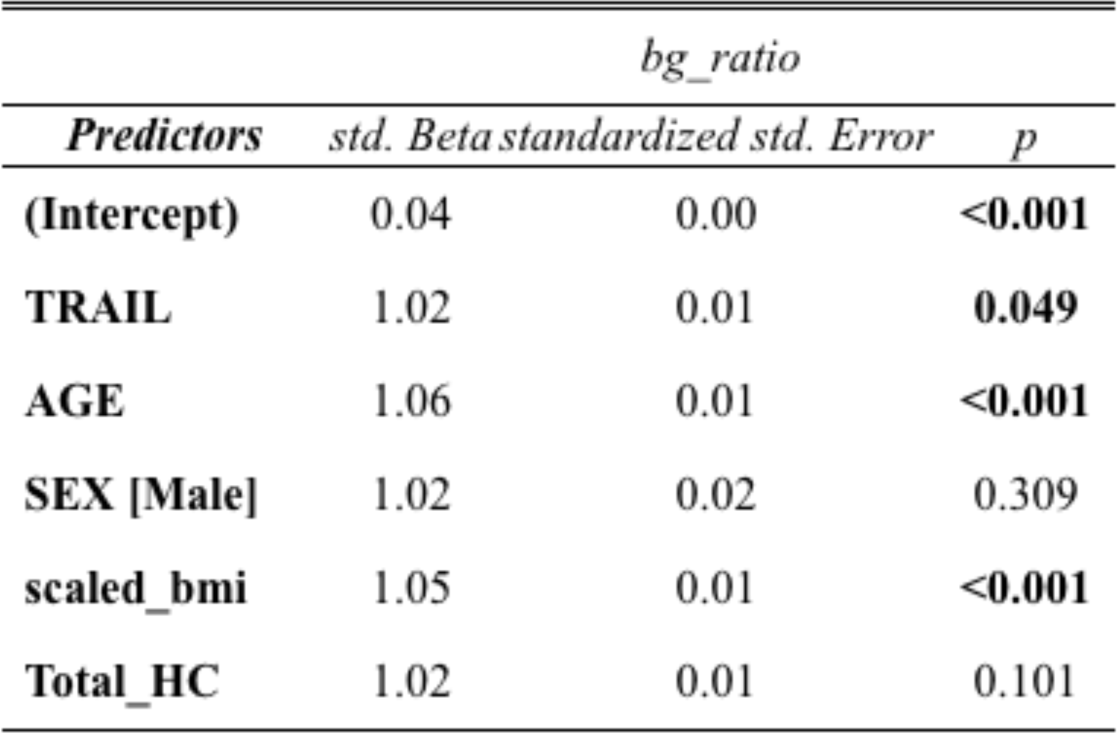
Beta coefficients, standard error and p-values for the main effect of TRAIL in the basal ganglia PVS volume fraction.

#### 3.2.3. Interaction analysis with Cortisol

There was no significant interaction between cortisol and any of the inflammatory biomarkers, except for CRP (p= 0.042, **Table 5**). Age and BMI were covariates that had significant direct associations. Statistical significance was confirmed when considering the medication use history (p=0.041), while a tendency of significance was seen with medication duration (p=0.053).

**Table 5:**
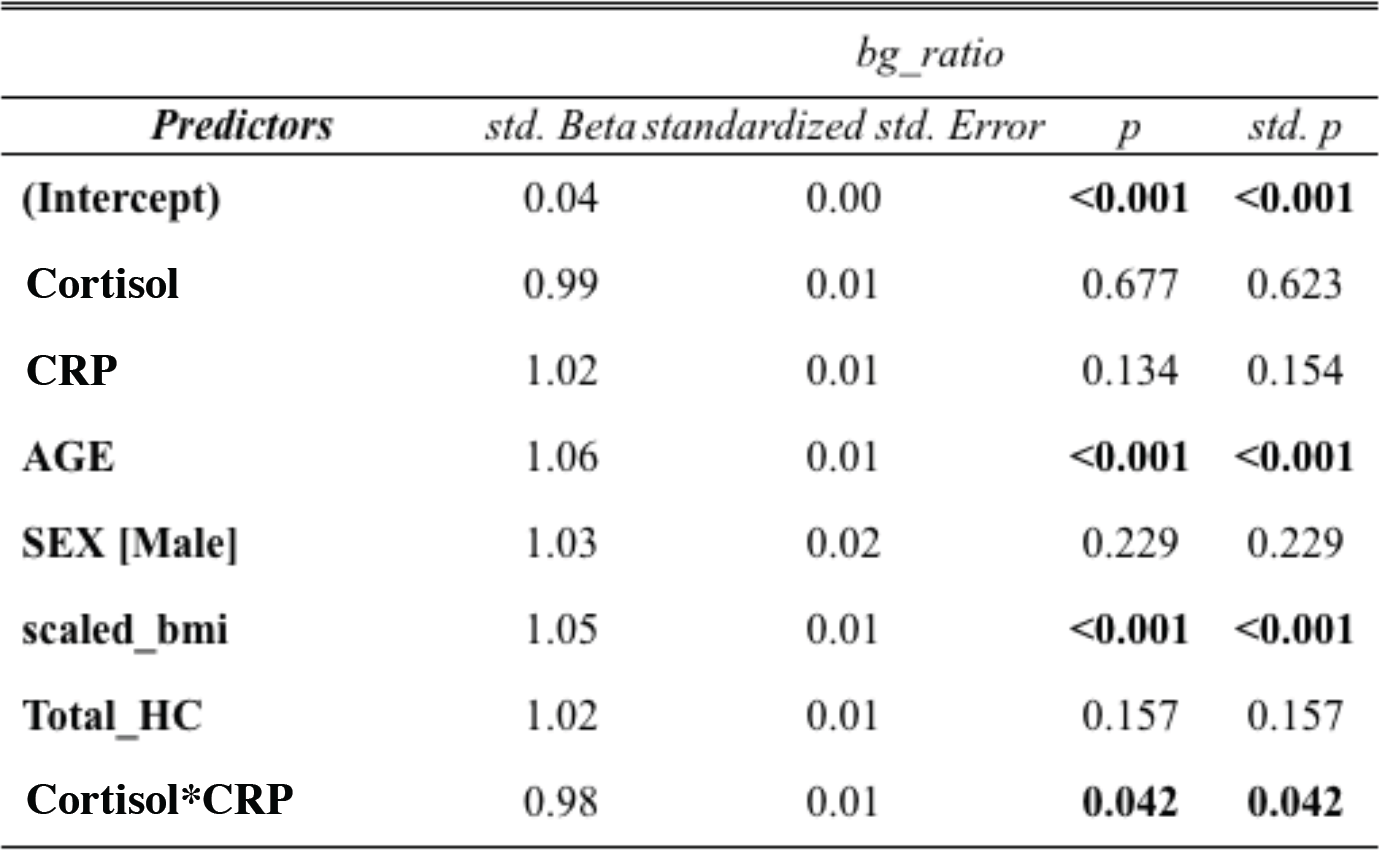
Beta coefficients, standard error and p-values for the interaction between Cortisol and CRP in the basal ganglia PVS volume fraction

#### 3.2.4. Interaction analysis with ACE

No significant interactions were shown between ACE and any of the inflammatory biomarkers.

#### 3.3. Sensitivity analysis of ApoE4

When we ran the analyses including ApoE4 as a model covariate representing genetic predisposition to Alzheimer’s disease, the presented results did not change. This means that adjusting for the number of ApoE4 allele copies does not alter our findings.

### 3.4. Within-group analysis based on cognitive group

To understand more the effect of cognitive impairment on the PVS volume fraction, we ran within-group analyses by dividing our sample in HC (n=48), MCI (n=325) and AD (n=92) groups. We ran the same regression models as for the entire population. Beta coefficients are shown in the tables in **Section 5** of Appendix.

#### 3.4.1. Healthy controls

Interaction of Cortisol with TNFr2 showed inverse associations with PVS (p= 0.037), such that higher levels of inflammation reduced Cortisol associations with PVS volume fraction in the basal ganglia. No other significant associations were seen.

#### 3.4.2. MCI

The MCI group represented the majority of our original sample (70%). Results for the CSO-PVS showed a significant negative main effect of TNFr2 (p= 0.023) and an inverse association with cortisol when interacting with TNF-*a* (p=0.008).

In the PVS of basal ganglia, a significant negative effect of TNF-*a* was seen (p=0.018), and a significant inverse association with ACE when interacting with MMP-2 (p=0.014).

#### 3.4.3. AD patients

In the AD group, interaction of Cortisol with MMP-9 showed inverse associations with CSO-PVS (p<0.001), such that higher levels of MMP-9 reduced Cortisol associations with PVS volume fraction. A positive effect of Cortisol was also shown (p=0.04).

In the basal ganglia, interaction of ACE with CRP showed a borderline inverse association with PVS volume fraction (p=0.048).

Results with diagnosis as covariate are reported in the Section 4 of Appendix. Forest plots reporting beta coefficients are shown in **Figure A5**.

## 4. Discussion

This paper shows for the first time how the combined action of stress-related, hypertension-related and inflammatory plasma biomarkers are associated with structural alterations of PVS. In this section, we discuss results from the 1.5T data.

As recently suggested (Zeng et al., 2022), the relationship between structural alteration of PVS, neuroinflammation and neurodegeneration can be described as a “vicious cycle”. The accumulation of inflammatory cellular components can cause an enlargement of PVS, leading to Aβ accumulation; at the same time, neurodegeneration can lead to neuroinflammatory cascade with a consequent enlargement of PVS, decreasing the glymphatic clearance activity.

### 4.1. Association with TRAIL

We found a borderline direct association between PVS and TRAIL, that is larger PVS volume fraction was related to higher TRAIL levels in basal ganglia. TRAIL, also called Apo2L, belongs to the TNF superfamily and it functions as an inducer of cellular apoptosis, after being released by activated microglia during neuroinflammation (Gao et al., 2020). A previous study using immunohistochemistry showed the lack of TRAIL molecules in the brain, rather the presence of TRAIL receptors on astrocytes, neurons and oligodendrocytes (Dörr et al., 2002). This may suggest that rather than having a role in the immune defense mechanisms in the brain, TRAIL receptors can be involved as effectors of neuroinflammation, showing apoptosis-blocking and apoptosis-mediating actions in brain autoimmune diseases via TRAIL upregulation operated by T-cells (Dörr et al., 2002). Since neurons, oligodendrocytes and astrocytes have TRAIL receptors, the interaction between T-cells and TRAIL receptors can be fatal for the cell. Excessive apoptosis has been found to be associated to Alzheimer’s disease, stroke and multiple sclerosis (Burgaletto et al., 2020); when there is inflammation occurring, T-cells can cross the BBB and release neurotoxins that aggravate the neurodegenerative processes. Upregulated TRAIL was found in the brain of AD patients when compared to healthy controls, and the highest concentration was located in proximity to the Aβ plaques (Uberti et al., 2004). TRAIL is responsible for neuronal loss in neurodegenerative diseases, and it is associated with cognitive decline in AD mouse models (Burgaletto et al., 2020). There are no many studies reporting the association of TRAIL and PVS volume fraction; during inflammation, PVS enlargements have been seen to be caused by the formation and deposition of PVS macrophages (PVM) within the vascular walls in response to an inflammation event; the PVMs plays a role in the cerebrovascular dysfunction and in the production of reactive oxygen species caused by the aggregating of Aβ, culminating in a hyper-production of TRAIL and cell death (Park et al., 2017).

### 4.2. TNFr2 and ACE

A significant negative main effect of TNFr2 was found on the CSO-PVS volume fraction, indicating that lower concentrations of TNFr2 were associated with higher PVS volume fraction. The interaction of TNFr2 with ACE was also significantly inverse, showing that the CSO-PVS relationship with ACE was becoming more negative with higher levels of TNFr2. Such results could indicate the neuroprotective effect of TNFr2 already shown by previous studies (Ortí-Casañ et al., 2022) (Dong et al., 2016), expressed by promoting the degradation of Aβ. Higher levels of TNFr2 help to counteract the neuroinflammatory state, promoting neuronal survival and myelination following a brain insult (Dong et al., 2016). A recent study showed that the lack of TNFr2 at cellular and physiological level corresponded to over-expression of inflammatory genes and increased expression of inflammatory cytokines, culminating in increased BBB permeability (Madsen et al., 2020).

### 4.3. Interactions with Cortisol

Results show a significant interaction between Cortisol and CRP for the PVS volume fraction in the basal ganglia. In the CSO-PVS, significant interactions were seen between Cortisol and TNF-α, CRP, MMP9, TNFr2 and CRP. These findings suggest that the association between Cortisol and PVS volume fraction varies across different levels of the inflammatory markers. This relationship is influenced by older age and higher BMI in older adults with cognitive impairment, previously shown to influence vascular contractility and to contribute to PVS enlargement (Barisano et al., 2020).

In the healthy brain, cortisol is a powerful anti-inflammatory hormone, limiting the spread of damaged cells in the brain, and regulating the levels of glucose (Hannibal & Bishop, 2014). The cortisol peak occurs in the morning and decreases during the day, and PVS volume fraction has been shown to be lower in the morning and higher in the afternoon (Barisano et al., 2020), suggesting that the hormonal concentration may be a factor influencing the changes of PVS volume fraction.

During a brain insult, activation of macrophages and microglia can release pro-inflammatory cytokines and lead to hyper-production of cortisol as a stress response factor, contributing to faster accumulation of Aβ and accelerated cognitive impairment (Toledo et al., 2012) (Pistollato et al., 2016).

Neuroinflammation has a positive effect on the brain in healthy conditions, for example it increases blood flow and removes phagocytosis. The microglia-macrophages system represents the key cells for the immune response, but they can also trigger an overproduction of cytokines, which lead to BBB disruption and alteration of neurogenesis in pathological conditions.

Interaction analyses showed that in the CSO, the relationship between PVS volume fraction with cortisol depends on the levels of inflammatory biomarkers (i.e., higher levels of inflammation reduced cortisol associations with PVS).

The dysregulation of glucocorticoids production (including cortisol) associated with aging has an impact on the inflammatory processes and the activation of the immune system (Valbuena Perez et al., 2020). In an experimental study, the levels of proinflammatory cytokines (in particular TNF-*a* and CRP) were higher in aged mice (Valbuena Perez et al., 2020), along with hypothalamic-pituitary-adrenal (HPA) axis dysregulations with a consequent reduction in glucocorticoids synthesis. In addition, CRP levels can differ across individuals as it was found to be influenced by lifestyle factors, for example obesity and smoking (Natale et al., 2022), and ethnicity (Morimoto et al., 2014) (Watanabe et al., 2016).

Another explanation of such findings is the connection between cognitive alterations and dysfunction in the negative feedback system of the HPA axis (de Souza-Talarico et al., 2011). This miscommunication leads to higher production of cortisol which has been seen in MCI and AD patients. Previous studies have also shown that individuals with cognitive impairment (starting from MCI stages onwards) present higher basal levels of cortisol compared to healthy old people, and this is due to increased and faster accumulation of Aβ in the brain (Pistollato et al., 2016) (Boespflug et al., 2018b), which has been associated with an enlarged PVS volume fraction (Sepehrband et al., 2021).

It has been proposed that cortisol plays a dual effect in regulating inflammation processes and immune response (Yeager et al., 2011). Glucocorticoids (GCs), whose cortisol is the most common one, have both a pro and anti-inflammatory effect. Levels of glucocorticoids expressed are key for an effective response during an acute systemic inflammation event: decreases of GCs trigger prolonged inflammation, while increased concentration of GC impairs inflammatory response against infection, with a slowdown in tissue repair. In humans, GCs low levels are associated with a weaker immune response and recurrent infections; this highlights the balance between glucocorticoids activity and inflammatory mediators’ action is necessary for immune mechanisms and the removal of pathogens (Cruz-Topete & Cidlowski, 2015).

### 4.4. PVS in clinical settings

Most of the results showed statistically significant associations and interactions on the CSO-PVS rather than the BG-PVS. While alterations in BG-PVS are more linked to hypertension and small vessel diseases, CSO-PVS structural alterations are associated with beta amyloid deposition. This was confirmed by a retrospective study (Kim et al., 2021), that found higher CSO-PVS linked to neurodegenerative processes and cognitive impairment, reflecting an increased deposition of Aβ in the brains of patients with Alzheimer’s disease. Another study looked at PVS changes in patients with intracerebral hemorrhage who developed cerebral amyloid angiopathy (CAA) in an Asian population (Tsai et al., 2021). They found that patients who developed CAA showed higher degree of CSO-PVS associated with higher vascular Aβ retention compared to those with low-degree CSO-PVS. High degree of CSO-PVS was also found in microglia-related inflammation in older people (Zeng et al., 2022), suggesting that inflammatory processes can lead to the dysfunction of the glymphatic system and deposition of Alzheimer’s disease-related biomarkers (i.e., Aβ and Tau).

The accumulation of Aβ and other neurotoxic at the level of PVS cause not only an enlargement at the structural level, but it also triggers an inflammatory accumulation of platelets which can lead to thrombotic events. Higher levels of CRP have been found to be related to release of pro-coagulant factors during neuroinflammation. CRP is generally considered a biomarker of acute inflammation, but it has also been shown to play a role in chronic inflammatory response (for example in neurodegeneration) (Noble et al., 2010). Lower levels of CRP have been associated to a faster cognitive impairment and found in patients with mild and moderate Alzheimer’s disease. CRP was also seen to be involved in developing cognitive decline in stroke patients, promoting Alzheimer’s disease progression after ischemia (Luan et al., 2018). CRP levels can stay stable over time and does not show variations during the day. This suggests CRP can be a stable and valid measure in clinical settings to investigate structural and functional alterations of the brain vasculature in neurodegenerative diseases.

### 4.5. Strengths and limitations

One of the novelties of this study is the automated segmentation of PVS applied on T1w images acquired at 1.5T. Image resolution is a key feature for the highest detection of PVS; the reason why we used 1.5T is that in the ADNI-1 cohort, most of the data were acquired at 1.5T and only a smaller sub-sample was available at 3T. The voxel resolution was higher in 1.5T than in 3T, and this also allowed us to compare the algorithm efficiency on data with different magnetic field strength. The automated technique of PVS segmentation represents a methodological strength for this study, as most of the literature on neuroinflammation and neuroimaging of PVS uses enlarged PVS based visual rating scores.

One of the limitations of this study is the lack of 3D FLAIR images in the ADNI-1 population, which is generally used to locate and remove WMHs. The decreased specificity of T1w images in respect to vascular abnormalities may lead to inaccurate detection of PVS; we investigated PVS changes in CSO and BG, as they are brain areas mostly associated with PVS in the literature. In this study, we included hippocampal volume as a sign of cognitive impairment and level of brain atrophy. The main results shown in this study considered the population as one group of older individuals, although their diagnoses included CN, MCI and AD. We ran further analyses by considering the diagnosis as covariate, as well as by splitting the population based on diagnosis; the sample size for each group was quite unbalanced (the majority of our participants were MCIs - 70% MCI, 20% AD and 10% CN-) y and this could affect the results. Another limitation is the lack of plasma Aβ and Tau measures in the analysis as markers of Alzheimer’s disease. This was not possible as there were missing data in more than half of participants in the ADNI-1 cohort. We did test for the APOE4 allele presence as a covariate in the model (coded based on how many copies of allele each participant had), which did not change our results.

## 5. Conclusion

This study shows how inflammation affects PVS volume fraction in older people with cognitive impairment, and how levels of stress and hypertension contribute to neuroinflammatory processes. The PVS volume fraction in the centrum semiovale showed significant inverse relationships with Cortisol in the presence of higher levels of inflammatory markers belonging to the TNF family, CRP and MMP-9. For the PVS in basal ganglia, we found a significant positive main effect of TRAIL, and a significant interaction term between Cortisol and CRP.

The concentration of cytokines is crucial for the physiological maintenance of the immune response in the brain; hyper-production can lead to accelerated accumulation of neurotoxins, decreasing the molecular waste clearance and leading to neuroimmune and neurodegenerative diseases. Future studies should build on these findings and look at the type of cells that are mostly found in the PVS during neuroinflammation, and how inflammatory markers can disrupt the fluid exchange that weaken the glymphatic system. Understanding if higher levels of plasma biomarkers lead to enlargement of PVS or vice versa can be helpful to develop therapeutic solutions to prevent neural death.

## Competing Interest Statement

The perivascular space mapping technology is part of a pending patent owned by JC. JC declares status as employee of the private company NeuroScope Inc (NY, USA).

## Acknowledgment

Authors would like to thank Ms. Caroline O’Driscoll for the illustration figure used in the paper to show the influence of inflammation on perivascular spaces. This work is supported by the National Institutes of Health (NIH) grants number R01AG070825, R01NS128486, and R41AG073024.

