## Supplementary Material for "Perivascular spaces in Alzheimer’s disease are associated with inflammatory, stress-related, and hypertension biomarkers"

### Appendix

#### Section 1: Methods

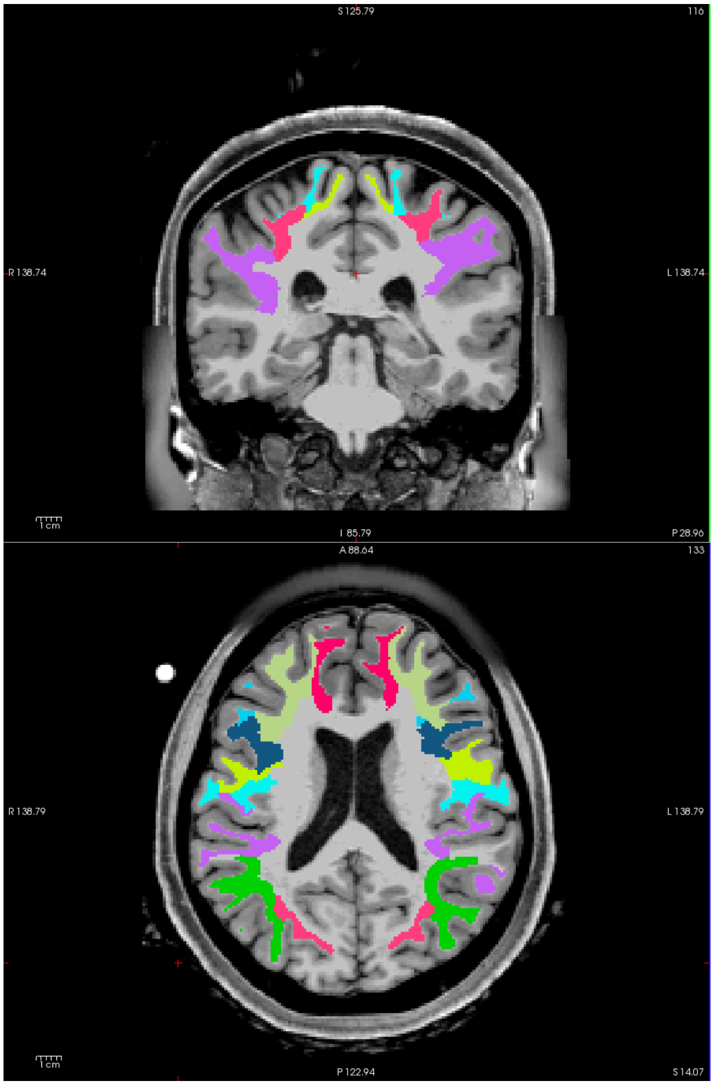

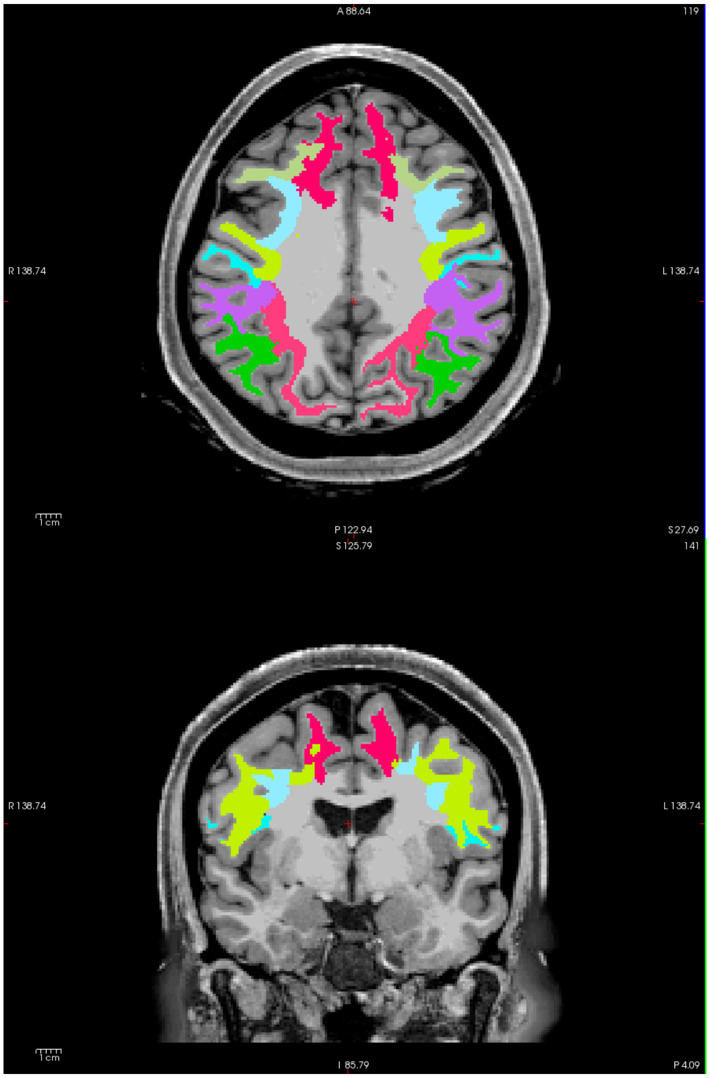


**Figure A1**: Centrum semiovale mask overlayed on T1w.

| **Labels** | **ROI Name** |
|---|---|
| 3003 | wm-lh-caudalmiddlefrontal |
| 3008 | wm-lh-inferiorparietal |

| | |
|---|---|
| 3018 | wm-lh-parsopercularis |
| 3019 | wm-lh-parsorbitalis |
| 3020 | wm-lh-parstriangularis |
| 3022 | wm-lh-postcentral |
| 3024 | wm-lh-precentral |
| 3027 | wm-lh-rostralmiddlefrontal |
| 3028 | wm-lh-superiorfrontal |
| 3029 | wm-lh-superiorparietal |
| 3031 | wm-lh-supramarginal |
| 4003 | wm-rh-caudalmiddlefrontal |
| 4008 | wm-rh-inferiorparietal |
| 4018 | wm-rh-parsopercularis |
| 4019 | wm-rh-parsorbitalis |
| 4020 | wm-rh-parstriangularis |
| 4022 | wm-rh-postcentral |

| | |
|---|---|
| 4024 | wm-rh-precentral |
| 4027 | wm-rh-rostralmiddlefrontal |
| 4028 | wm-rh-superiorfrontal |
| 4029 | wm-rh-superiorparietal |
| 4031 | wm-rh-supramarginal |

**Table A1.** Regions of interest (ROIs) to construct Centrum Semiovale region using FreeSurfer parcellation mask. The white matter labels are based on Desikan-Killiany-Tourville adult cortical parcellation atlas (Klein and Tourville, 2012). Left hemisphere denoted by `wm-lh` has 30XX added to the Label index, and the right hemisphere denoted by `wm-rh` has 40XX.

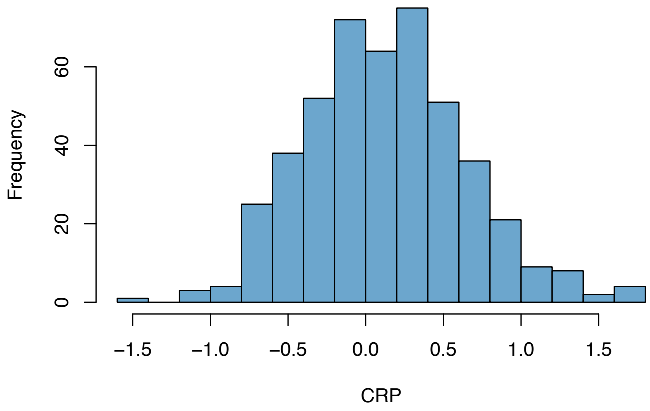


a)

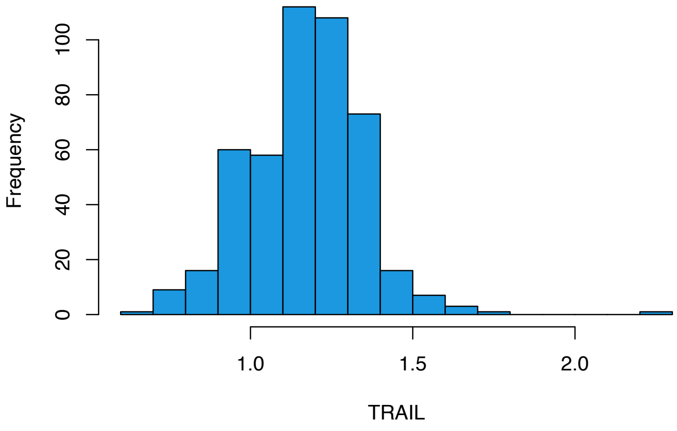
Frequency
0
20
40
60
80
100
1.0
1.5
2.0
TRAIL

b)

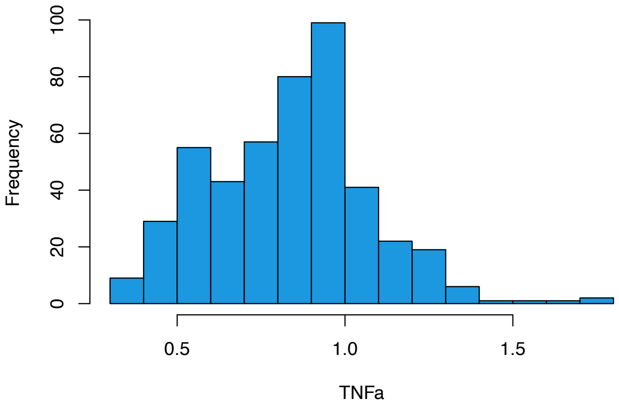
Frequency
0
20
40
60
80
100
0.5
1.0
1.5
TNFa

c)

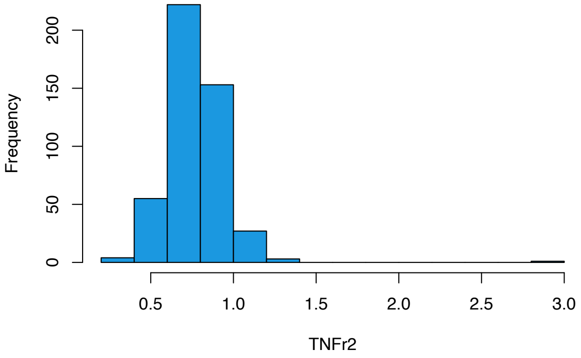
Frequency
0
50
100
150
200
0.5
1.0
1.5
2.0
2.5
3.0
TNFr2

d)

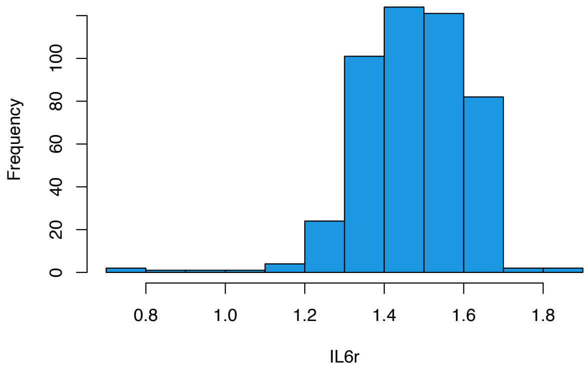


e)

**Figure A2**: Distribution of inflammatory biomarkers in the 1.5T cohort. a) CRP, b) TRAIL, c) TNFa, d) TNFr2, e) IL6r.

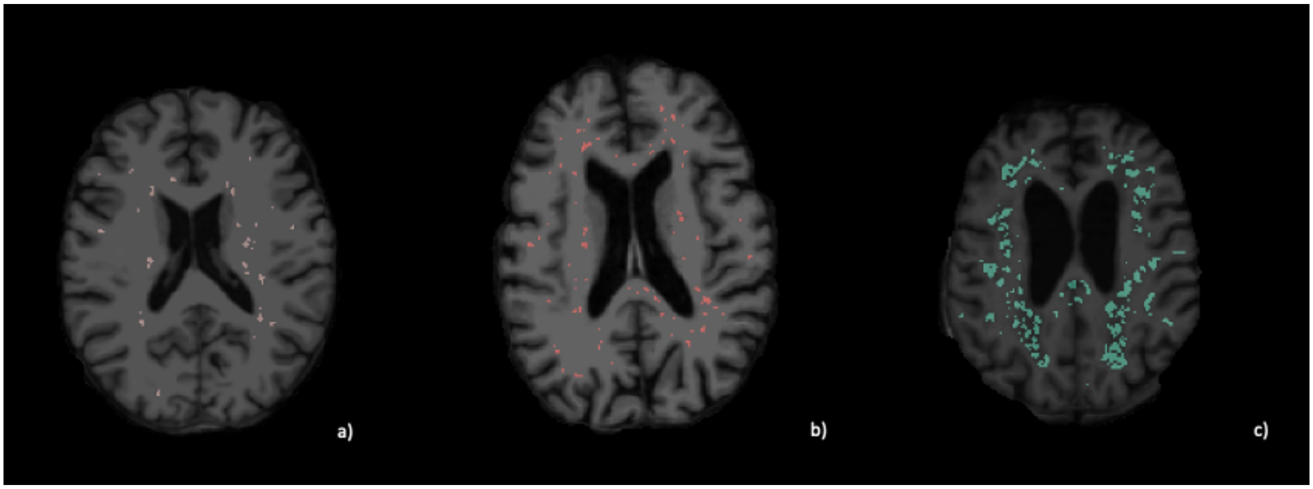


**Figure A3**: Visual representation of PVS segmentation in a healthy control (a), MCI (b), and AD patient (c).

#### Section 2: Results 3T

*Basal ganglia*

After removing the outliers, the final sample size was = 106. We ran the same model as the 1.5T data.

In the BG-PVS, a significant negative association was found with TRAIL (p=0.011, t =-2.578), with BMI and age being significant covariates.

TRAIL showed an independent effect also when we tested for the interaction with ACE and Cortisol; there was no significant interaction between ACE and TRAIL, but a significant effect for TRAIL was confirmed. Results were the same also when the medication history was added. A significant negative interaction was found also between MMP2 and ACE (p=0.0467, t=-2.014).

*Centrum semiovale*

In the centrum semiovale we did not find any significant association between PVS volume fraction and cortisol, ACE or any of the inflammatory markers.
Such difference between results in 3T and 1.5T might reflect how both the magnetic field strength and the sample size play a role in the association between PVS volume fraction and fluid biomarkers.

#### Section 3: Results 1.5T – negative results

a)

| | *CSO_ratio* | | | *CSO_ratio* | | | *CSO_ratio* | | | *CSO_ratio* | | |
|---|---|---|---|---|---|---|---|---|---|---|---|---|
| ***Predictors*** | *std. Beta* | *standardized std. Error* | *p* | *std. Beta* | *standardized std. Error* | *p* | *std. Beta* | *standardized std. Error* | *p* | *std. Beta* | *standardized std. Error* | *p* |
| **(Intercept)** | 0.02 | 0.00 | **<0.001** | 0.02 | 0.00 | **<0.001** | 0.02 | 0.00 | **<0.001** | 0.02 | 0.00 | **<0.001** |
| **Cortisol** | 1.02 | 0.03 | 0.453 | | | | | | | | | |
| **AGE** | 1.24 | 0.03 | **<0.001** | 1.24 | 0.03 | **<0.001** | 1.24 | 0.03 | **<0.001** | 1.24 | 0.03 | **<0.001** |
| **SEX [Male]** | 1.06 | 0.06 | 0.270 | 1.07 | 0.06 | 0.224 | 1.07 | 0.06 | 0.224 | 1.07 | 0.06 | 0.222 |
| **scaled_bmi** | 1.07 | 0.03 | **0.009** | 1.07 | 0.03 | **0.012** | 1.07 | 0.03 | **0.012** | 1.07 | 0.03 | **0.012** |
| **Total_HC** | 1.04 | 0.03 | 0.173 | 1.04 | 0.03 | 0.204 | 1.04 | 0.03 | 0.203 | 1.04 | 0.03 | 0.195 |
| **ACE** | | | | 1.00 | 0.03 | 0.971 | | | | | | |
| **MMP.2** | | | | | | | 1.00 | 0.03 | 0.992 | | | |
| **MMP.9** | | | | | | | | | | 1.01 | 0.03 | 0.690 |

b)

| | *bg_ratio* | | | *bg_ratio* | | | *bg_ratio* | | | *bg_ratio* | | |
|---|---|---|---|---|---|---|---|---|---|---|---|---|
| ***Predictors*** | *std. Beta* | *standardized std. Error* | *p* | *std. Beta* | *standardized std. Error* | *p* | *std. Beta* | *standardized std. Error* | *p* | *std. Beta* | *standardized std. Error* | *p* |
| **(Intercept)** | 0.04 | 0.00 | **<0.001** | 0.04 | 0.00 | **<0.001** | 0.04 | 0.00 | **<0.001** | 0.04 | 0.00 | **<0.001** |
| **Cortisol** | 0.99 | 0.01 | 0.608 | | | | | | | | | |
| **AGE** | 1.07 | 0.01 | **<0.001** | 1.07 | 0.01 | **<0.001** | 1.07 | 0.01 | **<0.001** | 1.07 | 0.01 | **<0.001** |
| **SEX [Male]** | 1.03 | 0.02 | 0.230 | 1.03 | 0.02 | 0.259 | 1.03 | 0.02 | 0.247 | 1.03 | 0.02 | 0.257 |
| **scaled_bmi** | 1.05 | 0.01 | **<0.001** | 1.05 | 0.01 | **<0.001** | 1.05 | 0.01 | **<0.001** | 1.05 | 0.01 | **<0.001** |
| **Total_HC** | 1.02 | 0.01 | 0.169 | 1.02 | 0.01 | 0.141 | 1.02 | 0.01 | 0.147 | 1.02 | 0.01 | 0.141 |
| **ACE** | | | | 1.01 | 0.01 | 0.404 | | | | | | |
| **MMP.2** | | | | | | | 1.00 | 0.01 | 0.830 | | | |
| **MMP.9** | | | | | | | | | | 1.00 | 0.01 | 0.737 |

**Figure A4:** Tables reporting beta coefficients and p-values for non-significant associations between stress-related (Cortisol), hypertension-related (ACE) and inflammatory-related (MMP-2 and MMP-9) biomarkers and PVS in CSO (a) and BG (b).

#### Section 4: Results from 1.5T with the diagnosis included as covariate in the model

Interaction of Cortisol with TNF-alpha (0.006), TNFr2 (p=0.003), TRAIL (p=0.018), MMP-9 (p=0.044) and CRP (p=0.015) showed inverse associations with PVS in the centrum semiovale.
Finally, interaction of Ace with TNFr2 (p=0.004) showed also inverse associations with PVS.

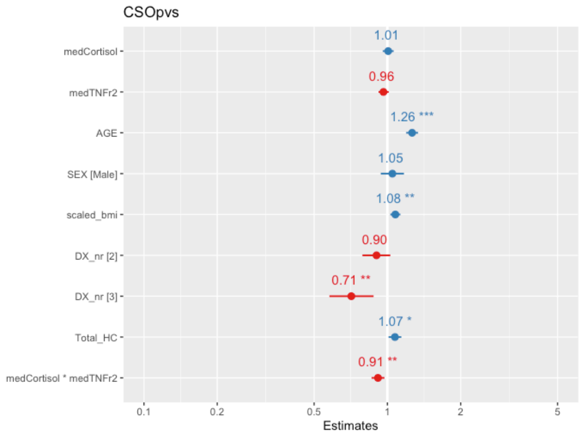


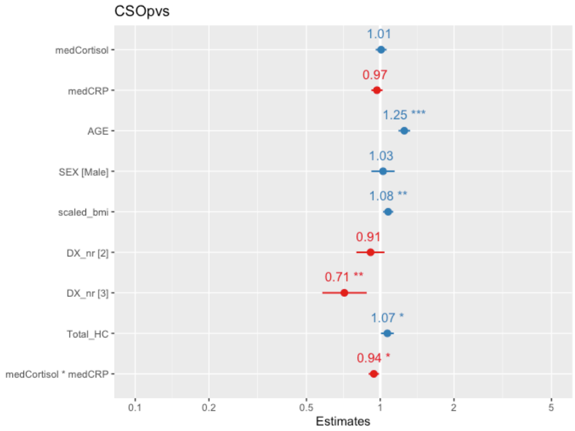


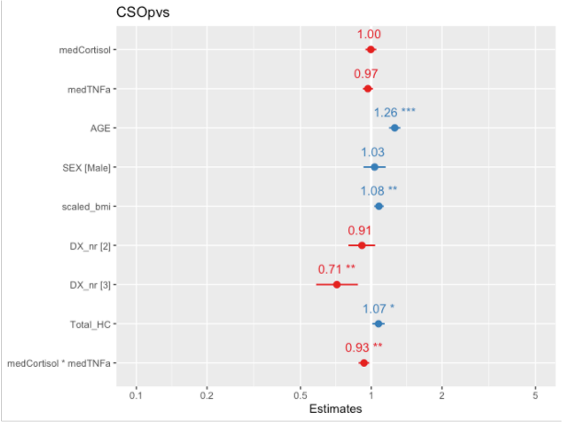


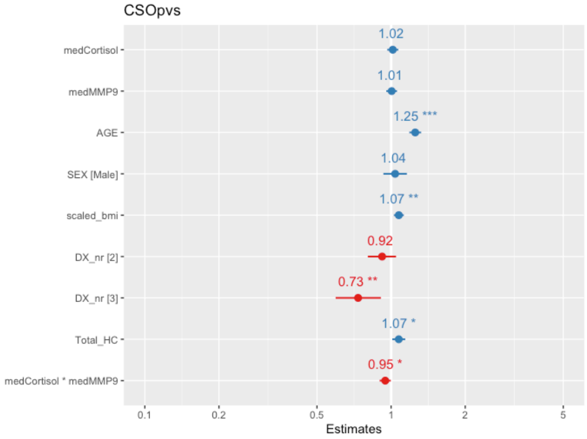


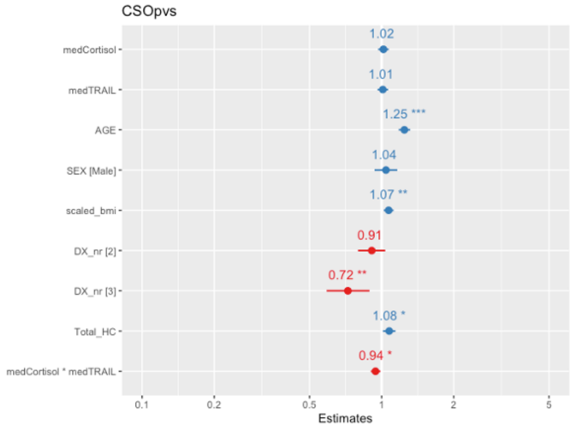


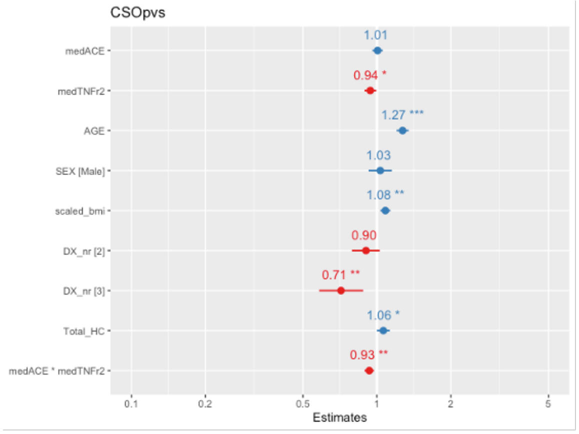

**Figure A5:** Forest plots representing the results of regression analyses including the diagnosis as covariate (categorized with 1= AD patients, 2= MCI and 3= healthy controls). The values in each plot represent standardized beta coefficients.

**Section 5**: Tables showing results of within-group analyses. Beta coefficients and p-values are reported in CN a), MCI b) and AD c).

| | *bg_ratio* | | | |
|---|---|---|---|---|
| ***Predictors*** | *std. Beta* | *standardized std. Error* | *p* | *std. p* |
| **(Intercept)** | 0.01 | 0.00 | **<0.001** | **<0.001** |
| **medCortisol** | 0.95 | 0.06 | 0.516 | 0.409 |
| **medTNFa** | 0.98 | 0.06 | 0.604 | 0.704 |
| **AGE** | 1.07 | 0.07 | 0.291 | 0.291 |
| **SEX [Male]** | 0.77 | 0.09 | **0.036** | **0.036** |
| **scaled_bmi** | 1.06 | 0.06 | 0.314 | 0.314 |
| **Total_HC** | 0.97 | 0.06 | 0.686 | 0.686 |
| **medCortisol * medTNFa** | 0.97 | 0.09 | 0.774 | 0.774 |

a) Healthy controls (CN) results. In the table standardized Beta coefficients, standard error and p-values are reported

| | *ratio_CS* | | | |
|---|---|---|---|---|
| ***Predictors*** | *std. Beta* | *standardized std. Error* | *p* | *std. p* |
| **(Intercept)** | 0.01 | 0.00 | **<0.001** | **<0.001** |
| **medCortisol** | 1.03 | 0.04 | 0.929 | 0.506 |
| **medTNFa** | 0.94 | 0.04 | 0.383 | 0.170 |
| **AGE** | 1.26 | 0.06 | **<0.001** | **<0.001** |
| **SEX [Male]** | 1.08 | 0.10 | 0.374 | 0.374 |
| **scaled_bmi** | 1.06 | 0.04 | 0.158 | 0.158 |
| **Total_HC** | 1.08 | 0.05 | 0.104 | 0.104 |
| **medCortisol * medTNFa** | 0.90 | 0.04 | **0.009** | **0.009** |

| | *ratio_CS* | | |
|---|---|---|---|
| ***Predictors*** | *std. Beta* | *standardized std. Error* | *p* |
| **(Intercept)** | 0.01 | 0.00 | **<0.001** |
| **TNFr.2** | 0.90 | 0.04 | **0.023** |
| **AGE** | 1.31 | 0.06 | **<0.001** |
| **SEX [Male]** | 1.12 | 0.10 | 0.229 |
| **scaled_bmi** | 1.08 | 0.05 | 0.069 |
| **Total_HC** | 1.05 | 0.05 | 0.259 |

| | *bg_ratio* | | | |
|---|---|---|---|---|
| ***Predictors*** | *std. Beta* | *standardized std. Error* | *p* | *std. p* |
| **(Intercept)** | 0.01 | 0.00 | **<0.001** | **<0.001** |
| **medACE** | 1.04 | 0.02 | 0.107 | **0.043** |
| **medMMP2** | 0.96 | 0.02 | 0.114 | 0.093 |
| **AGE** | 1.15 | 0.03 | **<0.001** | **<0.001** |
| **SEX [Male]** | 1.04 | 0.05 | 0.401 | 0.401 |
| **scaled_bmi** | 1.05 | 0.02 | **0.032** | **0.032** |
| **Total_HC** | 1.05 | 0.02 | **0.046** | **0.046** |
| **medACE * medMMP2** | 0.91 | 0.03 | **0.014** | **0.014** |

| | *bg_ratio* | | |
|---|---|---|---|
| ***Predictors*** | *std. Beta* | *standardized std. Error* | *p* |
| **(Intercept)** | 0.01 | 0.00 | **<0.001** |
| **TNF.alpha** | 0.95 | 0.02 | **0.018** |
| **AGE** | 1.15 | 0.03 | **<0.001** |
| **SEX [Male]** | 1.05 | 0.05 | 0.324 |
| **scaled_bmi** | 1.05 | 0.02 | **0.014** |
| **Total_HC** | 1.05 | 0.03 | 0.055 |

b) Results for the MCI group. In the table standardized Beta coefficients, standard error and p-values are reported

| | *bg_ratio* | | | |
|---|---|---|---|---|
| ***Predictors*** | *std. Beta* | *standardized std. Error* | *p* | *std. p* |
| **(Intercept)** | 0.01 | 0.00 | **<0.001** | **<0.001** |
| **medACE** | 1.01 | 0.04 | 0.810 | 0.744 |
| **medCRP** | 1.04 | 0.04 | 0.298 | 0.373 |
| **AGE** | 1.16 | 0.05 | **0.001** | **0.001** |
| **SEX [Male]** | 1.06 | 0.09 | 0.483 | 0.483 |
| **scaled_bmi** | 1.14 | 0.05 | **0.002** | **0.002** |
| **Total_HC** | 1.01 | 0.05 | 0.819 | 0.819 |
| **medACE * medCRP** | 0.92 | 0.04 | **0.048** | **0.048** |

| | *ratio_CS* | | | |
|---|---|---|---|---|
| ***Predictors*** | *std. Beta* | *standardized std. Error* | *p* | *std. p* |
| **(Intercept)** | 0.01 | 0.00 | **<0.001** | **<0.001** |
| **medCortisol** | 1.11 | 0.10 | **0.040** | 0.239 |
| **medMMP9** | 1.01 | 0.09 | 0.398 | 0.924 |
| **AGE** | 1.58 | 0.14 | **<0.001** | **<0.001** |
| **SEX [Male]** | 1.26 | 0.24 | 0.219 | 0.219 |
| **scaled_bmi** | 1.21 | 0.10 | **0.027** | **0.027** |
| **Total_HC** | 1.22 | 0.12 | **0.043** | **0.043** |
| **medCortisol * medMMP9** | 0.73 | 0.06 | **<0.001** | **<0.001** |

c) Results for AD group. In the table standardized Beta coefficients, standard error and p-values are reported.
